# Application of database-independent approach to assess the quality of OTU picking methods

**DOI:** 10.1101/042812

**Authors:** Patrick D. Schloss

## Abstract

Assigning 16S rRNA gene sequences to operational taxonomic units (OTUs) allows microbial ecologists to overcome the inconsistencies and biases within bacterial taxonomy and provides a strategy for clustering similar sequences that do not have representatives in a reference database. I have applied the Matthew’s correlation coefficient to assess the ability of 15 reference-independent and ‐dependent clustering algorithms to assign sequences to OTUs. This metric quantifies the ability of an algorithm to reflect the relationships between sequences without the use of a reference and can be applied to any dataset or method. The most consistently robust method was the average neighbor algorithm; however, for some datasets other algorithms matched its performance.

---

Numerous algorithms have been developed for solving the seemingly simple problem of assigning 16S rRNA gene sequences to operational taxonomic units (OTUs). These algorithms were recently the subject of benchmarking studies performed by Westcott and myself (1, 2), He et al (3), and Kopylova et al (4). These studies provide a thorough review of the sequencing clustering landscape, which can be divided into three general approaches: (i) *de novo* clustering where sequences are clustered without first mapping sequences to a reference database, (ii) closed-reference clustering where sequences are clustered based on the references that the sequences map to, and (iii) open reference clustering where sequences that do not map adequately to the reference are then clustered using a *de novo* approach. Assessing the quality of the clustering assignments has been a persistent problem in the development of clustering algorithms.

The recent analysis of Kopylova et al (4) repeated many of the benchmarking strategies employed by previous researchers. Many algorithm developers have clustered sequences from simulated communities or sequencing data from synthetic communities of cultured organisms and quantified how well the OTU assignments matched the organisms’ taxonomy (5–16). Although an OTU definition would ideally match bacterial taxonomy, bacterial taxonomy has proven itself to be fluid and reflect the biases of various research interests. Furthermore, it is unclear how the methods scale to sequences from the novel organisms we are likely to encounter in deep sequencing surveys. In a second approach, developers have compared the time and memory required to cluster sequences in a dataset (6, 13, 17, 18). These are valid parameters to assess when judging a clustering method, but indicate little regarding the quality of the OTU assignments. For example, reference-based methods are very efficient, but do a poor job of reflecting the genetic diversity within the community when novel sequences are encountered (2). In a third approach, developers have compared the number of OTUs generated by various methods for a common dataset (4, 5). Although methods need to guard against excessive splitting of sequences across OTUs, by focusing on minimizing the number of OTUs in a community developers risk excessively lumping sequences together that are not similar. In a fourth approach, a metric of OTU stability has been proposed as a way to assess algorithms (3). Although it is important that the methods generate reproducible OTU assignments when the initial order of the sequences is randomized, this metric ignores the possibility that the variation in assignments may be equally robust or that the assignments by a highly reproducible algorithm may be quite poor. In a final approach, some developers have assessed the quality of clustering based on the method’s ability to generate the same OTUs generated by other methods (18, 19). Unfortunately, without the ability to ground truth any method, such comparisons are tenuous. There is a need for an objective metric to assess the quality of OTU assignments.

Westcott and I have proposed an unbiased and objective method for assessing the quality of OTU assignments that can be applied to any collection of sequences (1, 2). Our approach uses the observed dissimilarity between pairs of sequences and information about whether sequences were clustered together to quantify how well similar sequences are clustered together and dissimilar sequences are clustered apart. To quantify the correlation between the observed and expected OTU assignments, we synthesize the relationship between OTU assignments and the distances between sequences using the Matthew’s correlation coefficient (20). I have expanded our previous analysis to evaluate three hierarchical and seven greedy de *novo* algorithms, one open-reference clustering algorithm, and four closed-reference algorithms (Figure 1). To test these approaches I applied each of them to datasets from soil (21), mouse feces (22), and two simulated datasets. The simulated communities were generated by randomly selecting 10,000 16S rRNA sequences that were unique within the V4 region from the SILVA non-redundant database (4, 23). Next, an even community was generated by specifying that each sequence had a frequency of 100 reads and a staggered community was generated by specifying that the abundance of each sequence was a randomly drawn a uniform distribution between 1 and 200. A reproducible version of this manuscript and analysis has been added to the repository available at https://github.com/SchlossLab/Schloss_Cluster_PeerJ_2015.

**Figure 1.**
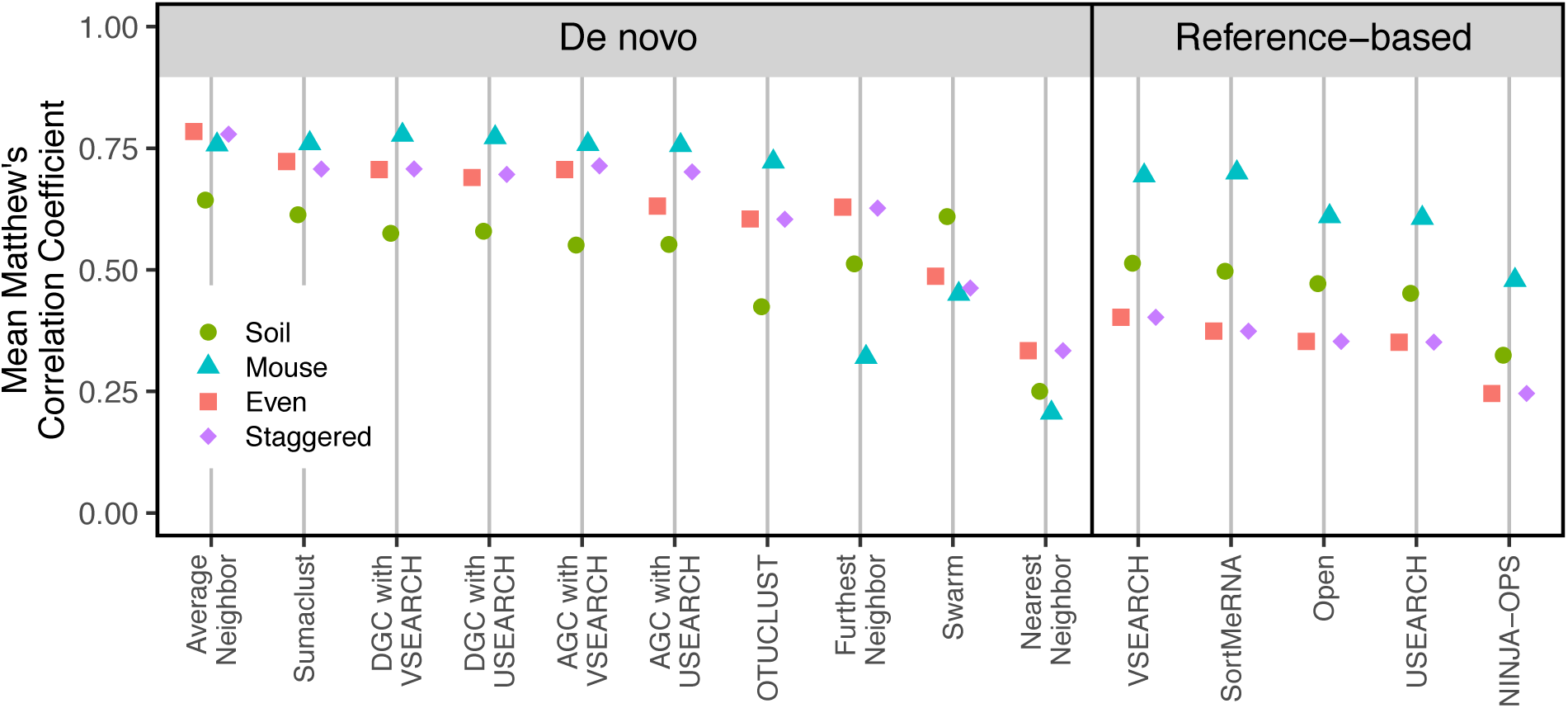
Comparison of OTU quality generated by multiple algorithms applied to four datasets. The nearest, average, and furthest neighbor clustering algorithms were used as implemented in mothur (v.1.37)(24). Abundance (AGC) and Distance-based greedy clustering (DGC) were implemented using USEARCH (v.6.1) and VSEARCH (v.1.5.0)(3, 5, 25). Other *de novo* clustering algorithms included Swarm (v.2.1.1)(6, 7), OTUCLUST (v.0.1)(26), and Sumaclust (v.1.0.20). The MCC values for Swarm were determined by selecting the distance threshold that generated the maximum MCC value for each dataset. The USEARCH and SortMeRNA (v.2.0) closed-reference clusterings were performed using QIIME (v.1.9.1) (27, 28). Closed-reference clustering was also performed using VSEARCH (v.1.5.0) and NINJA-OPS (v.1.5.0) (16). The order of the sequences in each dataset was randomized thirty times and the intra-method range in MCC values was smaller than the plotting symbol. MCC values were calculated using mothur.

I replicated the benchmarking approach that I have used previously to assess the ability of an algorithm to correctly group sequences that are similar to each other and split sequences that are dissimilar to each other using the MCC (1, 2). When I compared the MCC values calculated using the ten *de novo* algorithms with the four datasets, the average neighbor algorithm reliably performed as well or better than the other methods (Figure 1). For the murine dataset, the MCC values for the VSEARCH (AGC: 0.76 and DGC: 0.78) and USEARCH-based (AGC: 0.76 and DGC: 0.77) algorithms, Sumaclust (0.76), and average neighbor (0.76) were similarly high. For each of the other datasets, the MCC value for the average neighbor algorithm was at least 5% higher than the next best method. Swarm does not use a traditional distance-based criteria to cluster sequences into OTUs and instead looks for natural subnetworks in the data. When I used the distance threshold that gave the best MCC value for the Swarm data, the MCC values were generally not as high as they were using the average neighbor algorithm. The one exception was for the soil dataset. Among the reference-based methods, all of the MCC values suffer because when sequences that are at least 97% similar to a reference are pooled, the sequences within an OTU could be as much as 6% different from each other. The effect of this is observed in the MCC values that were calculated for the OTUs assigned by these methods generally being lower than those observed using the *de novo* approaches (Figure 1). It is also important to note that the MCC values for the closed-reference OTUs are inflated because sequences were removed from the analysis if there was not a reference sequence that was more than 97% similar to the sequence. By choosing to focus on the ability to regenerate taxonomic clusterings, minimizing the number of OTUs, and performance Kopylova et al (4) concluded that Swarm and SUMACLUST had the most consistent performance among the *de novo* methods. My objective MCC-based approach found that Sumaclust performed well, but was matched or outperformed by the average neighbor algorithm; using a 3% threshold, Swarm was actually one of the worst methods. Given the consistent quality of the clusterings formed by the average neighbor algorithm, these results confirm the conclusion from the previous analysis that researchers should use the average neighbor algorithm or calculate MCC values for several methods and use the clustering that gives the best MCC value (2).

Next, I investigated the ability of the reference-based methods to properly assign sequences to OTUs. The full-length 16S rRNA gene sequences in the default reference taxonomy that accompanies QIIME are less than 97% similar to each other. Within the V4 region, however, many of the sequences were more similar to each other and even identical to each other. As a result, we previously found that there was a dependence between the ordering of sequences in the reference database and the OTU assignments with USEARCH and VSEARCH (2). To explore this further, we analyzed the 32,106 unique sequences from the murine dataset with randomized databases. VSEARCH always found matches for 27,737 murine sequences, the reference matched to those sequences differed between randomizations. For USEARCH there were between 28,007 and 28,111 matches depending on the order of the reference. In the updated analysis we found that SortMeRNA resulted in between 23,912 and 28,464 matches. Using NINJA-OPS with different orderings of the reference sequences generated the same 28,499 matches. These results point to an additional problem with closed-reference clustering, which is the inability for the method to assign sequences to OTUs when a similar reference sequence does not exist in the database. For the well-characterized murine microbiota, NINJA-OPS did the best by finding relatives for 88.8% of the unique murine sequences. As indicated by the variation in the number of sequences that matched a reference sequence, these methods varied in their sensitivity and specificity to find the best reference sequence. Of the closed-reference methods, NINJA-OPS had the best sensitivity (99.7%) and specificity (79.7%) while SortMeRNA had the worst sensitivity (95.7%) and VSEARCH had the worst specificity (60.3%). Reference-based clustering algorithms are much faster than *de novo* approaches, but do not generate OTUs that are as robust.

Although the goal of Kopylova et al (4) was to compare various clustering algorithms, they also studied these algorithms in the broader context of raw sequence processing, screening for chimeras, and removal of singletons. Each of these are critical decisions in a comprehensive pipeline. By including these steps, they confounded their analysis of how best to cluster sequences into OTUs. The effect of differences in MCC values on one’s ability to draw inferences is unclear and admittedly may be relatively minor for some datasets. Because of this uncertainty, researchers should use the most reliable methods available in case the differences in clustering do effect the conclusions that can be drawn from a particular dataset. Through the use of objective criteria that measure the quality of the clusterings, independent of taxonomy or database, researchers will be able to evaluate which clustering algorithm is the best fit for their data.

